# Neuromorphic Imaging Cytometry on Human Blood Cells

**DOI:** 10.1101/2024.11.16.623904

**Authors:** Ziyao Zhang, Haoxiang Yang, Jiayin Li, Shin Wei Chong, Jason K. Eshraghian, Ken-Tye Yong, Daniele Vigolo, Helen M. McGuire, Omid Kavehei

**Affiliations:** School of Biomedical Engineering, The University of Sydney, NSW, Australia; Faculty of Science, University of Technology Sydney, NSW, Australia; Department of Electrical and Computer Engineering, The University of California, Santa Cruz, USA; The University of Sydney Nano Institute, The University of Sydney, Sydney, NSW, Australia; School of Medical Science, The University of Sydney, NSW, Australia

**Keywords:** neuromorphic cytometry, neuromorphic sensor, event camera, event-based cytometry, image-based cytometry, image-enabled cell sorting, human blood cells

## Abstract

Image-enhanced cytometry and sorting are powerful technologies that provide single-cell resolution and, where possible, cell actuation based on spatial and fluorescence characterisation. With the emergence of deep learning (DL), numerous cytometry-related works incorporate DL to assist their research in handling data-intensive and repetitive workloads. The rich spatial information provided by single-cell images has exceptional use with DL models to classify cells, detect rare cell events, disclose irregularity and achieve higher sample purity than a conventional feature-gating strategy. One of the significant challenges in these image-enable technologies is the constrained throughput owing to the data-expensive image acquisition and balancing between speed and resolution. This work introduces a novel paradigm by adopting a bio-inspired neuromorphic photosensor to capture fast-moving cell events. It facilitates a data-efficient, fluorescence-sensitive, fast inference approach to establish a foundation for neuromorphic-enabled cytometry/sorting applications. We have also curated the first neuromorphic-encoded cell dataset, including human blood cells (red blood cells, neutrophils, lymphocytes, thrombocytes), endothelial cells and polystyrene-based microparticles. To evaluate the data quality and potential of DL-based gating, we have directly trained a hybrid classification model based on this dataset, accomplishing a promising performance of 97% accuracy and F1 score with a significant reduction in memory usage and power consumption. Combining neuromorphic imaging and DL holds substantial potential to develop into a next-generation AI-assisted cytometry and sorting application.

## 1 Introduction

Imaging flow cytometry (IFC) and image-enabled cell sorting (ICS) are indispensable technologies that exploit single-cell resolution and fluorescence intensity to quantitatively characterise cell phenotypes and provide isolation where possible.^1,2^ Such paradigm permits cells to be analysed with a detailed resolution of cellular composition, morphology and protein distribution.^3^ Both technologies offer visualisation on cells conventionally captured by frame-based sensors (FBSs) such as charged-coupled devices or complementary metal-oxide semiconductors.^4^ With ICS, cell sorting can be actuated based on collected spatial and fluorescence information. However, the throughput of these image-based technologies is inherently constrained by the data-extensive image acquisition and the trade-off between speed and resolution.^1,5,6^ FBSs with long frame intervals may fail to accommodate objects travelling at a higher velocity, leading to image degradation, misdetection and other motion-induced artefacts. Furthermore, characterising cells with frame-based feedback can accumulate enormous pixel data registered with blank and redundant background information that is irrelevant to cell events. Such data structures can be transfer-expensive in terms of storage and processing, especially considering the quantity of a cell population or populations. Thus, a pixel-efficient and high-temporal-resolution image acquisition method should be explored to overcome the traditional constraints of IFC and ICS.

Feature gating adopted in IFC and ICS often required manual intervention to classify or retrieve discrete populations using a series of 2D scatter plots that decompose morphological characteristics and fluorescence intensity.^7^ While robust, this strategy relies strongly on domain expertise and can be labour-intensive for large-scale experiments. Traditional gating of the high-dimensional data can limit the interpretation into bivariate hierarchical analysis of 2D scatter plots. However, these image-enabled systems were designed to offer multivariate spatial feedback to reveal complicated data structures. Moreover, cells being analysed are often labelled with multiple fluorescence tags for unique identification and evaluation. This procedure can be cost-expensive and compromise the viability and re-usability of cells for downstream applications such as regenerative medicine, cell-based therapy and precision medicine.^7,8^ In recent years, numerous computational endeavours have been implemented to automate cytometry data analysis for quality control, data visualisation and sample classification.^7^ The research conducted by Hayashi et al.^9^ reported a significant reduction in false-positive rates and an 11.0-fold sample enrichment using a DL classifier for ICS.

The neuromorphic vision sensor (NVS), also known as the event camera, dynamic vision sensor or silicon retina, is a bio-inspired photosensor that imitates the continuous signalling and key attention to motion information of biological retina.^10,11,12,13^ NVS can detect relative contrast changes in a scene with asynchronous firing pixels that transmit only the activated array address. This event-driven perception enables superior data efficiency, spatiotemporal resolution and dynamic range with low power consumption.^11,14^ In the context of IFC and ICS, NVS can report an efficient pixel collection that only focuses on cell phenotype instead of shuttering all pixels in a sensor for full-frame readout. Our previous attempts conceptualised a neuromorphic imaging cytometry/sorting (NICS) to overcome the traditional frame-based challenges, demonstrating a superior fluorescence sensitivity in detecting fluorescence microscale targets.^15,16,17^

To the best of our knowledge, we present the first label-free neuromorphic cell dataset including human blood cells (red blood cells, neutrophils, lymphocytes, platelets), primary human umbilical vein endothelial cells (HUVECs) and polystyrene-based microparticles. This is the first neuromorphic biological cell dataset that contains a variety and quantity of microscale objects. Furthermore, to evaluate the image quality of the cell data and the plausibility of automating feature gating with a DL approach, we have developed and directly trained a hybrid model combining the convolutional block attention module (CBAM) with spiking neural network (SNN), referred to CBAM-SNN for validation. This model has achieved a 97% accuracy and F1 score with a significant reduction in memory usage and power consumption compared to a classic convolutional neural network (CNN). Such an architecture shows much potential for feature gating and fast inference in NICS-like applications. We believe that NICS holds great potential for overcoming the everlasting FBS constraints in IFC and ICS, developing into a next-generation AI-assisted cytometry/sorting application.

## 2 Results & Discussion

### 2.1 Neuromorphic imaging platform

Human blood samples and other sample cells undergo bulk separation to obtain targeted cell lines. Cells are introduced into our developed neuromorphic imaging platform, as shown in Fig. 1. As the cells pass through the view of interest, an NVS records the cell features with their respective pixel address, timestamp and polarity (X, Y, P, T). Such information can be divided into two usages: (1) Collected events are transformed into tensors and then inferred by the developed CBAM-SNN to conclude its respective class. (2) Events are transformed into frame-based outputs to examine the integrity of captured features and allow calibration work to optimise the visual effects. Then, these data are curated into the dataset for further training and model development purposes. The specifications of the EVS and other sensors used in ICS, cell tracking or fluorescence detection were collected in Table 1, highlighting a superior temporal resolution, dynamic range and power consumption of EVS.

**Table 1:** Comparison of event camera (EVK4) with other cameras used in cell imaging or sorting works for either cell detection, trajectory monitoring or fluorescence detection. The frame rates recorded below are at the sensors’ full resolution and can be increased with reduced dimensions.

|  | Current work | Nawaz et al<sup>2,23</sup> | Lee et al<sup>24</sup> | Isozaki et al<sup>25</sup> |
| --- | --- | --- | --- | --- |
| Vision Sensor | EVK4 (IMX636) | EoSens CL (MC1362) | DMK 37AUX287 | Phantom v2640 |
| Resolution (Pixels) | 1280 × 720 | 1280 × 1024 | 720 × 540 | 2048 × 1952 |
| Equivalent Frame Rate (fps) | > 10,000 | 500 | 539 | 4,855 |
| Pixel Size (μm) | 4.86 | 14 | 6.9 | 13.5 |
| Dynamic Range (dB) | > 120 | 60 | 60 | 64.1 |
| Power Consumption (W) | 0.5 ~ 1.5 | 7 | 1.7 | 280 |
| Output | events | images/frames | images/frames | images/frames |

**Figure 1.**
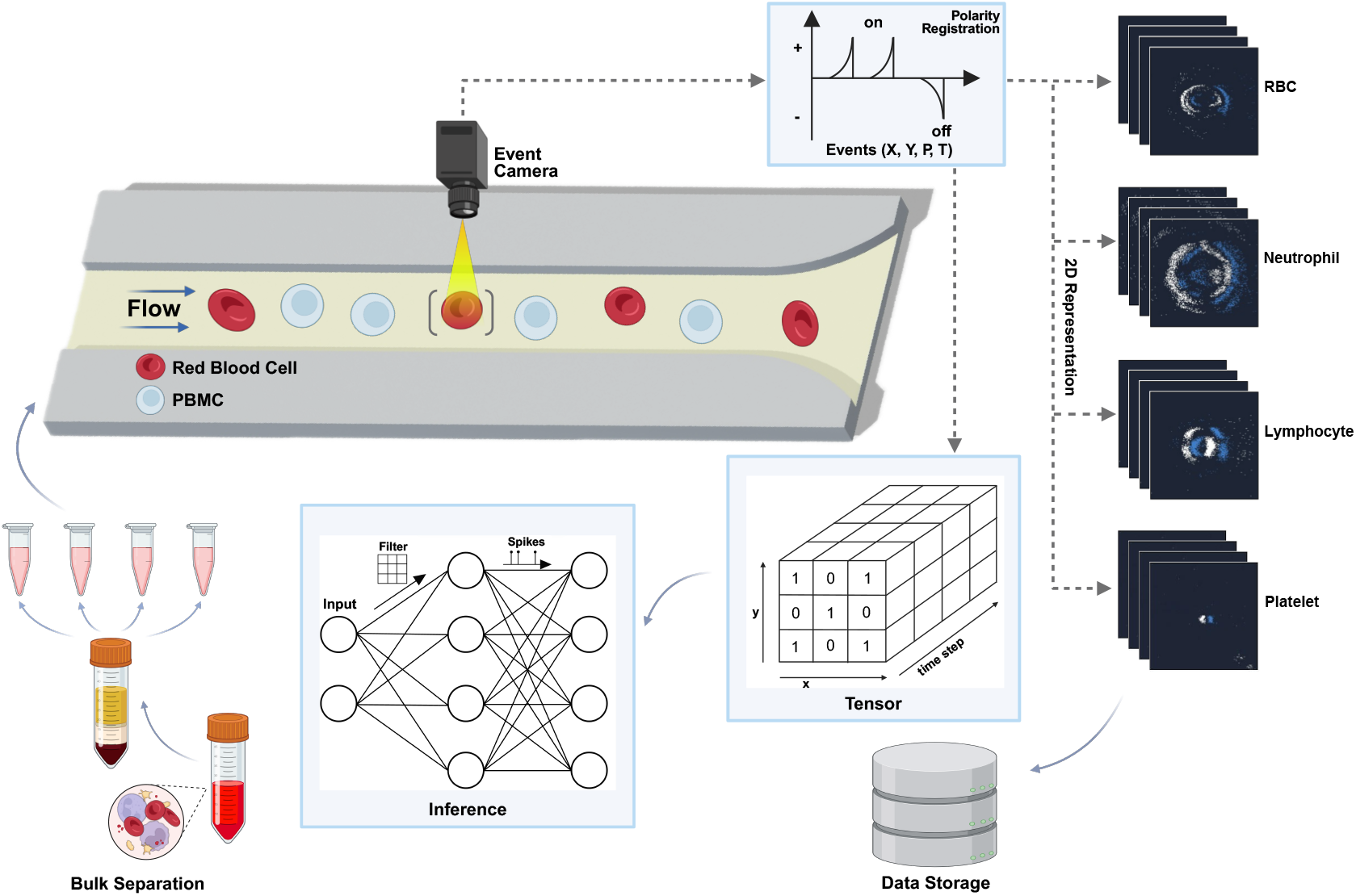
Schema of neuromorphic imaging platform. The samples were prepared by bulk separation to extract the cells of interest. As the cells were introduced into the microfluidic channel, the cell events were registered by EVS for two usages: **(1)** cell events were transformed into 2D images for visualisation and data curation. **(2)** Cell events were transformed into tensors that undergo a neural network for cell classification.

### 2.2 Data curation

Red blood cells (RBCs), neutrophils, lymphocytes, platelets, HUVECs and 8-*μ*m particles were curated in an event format and image format at a resolution of 224 × 224 pixels for illustration in Fig. 2. Cell images containing multiple targets were included for enhanced generalisation and robustness. 8 *μ*m particles were added to resemble similar sizes of RBCs and lymphocytes to potentially increase the data complexity and constrain the model for low-dimensional feature extraction by only size variants.

**Figure 2.**
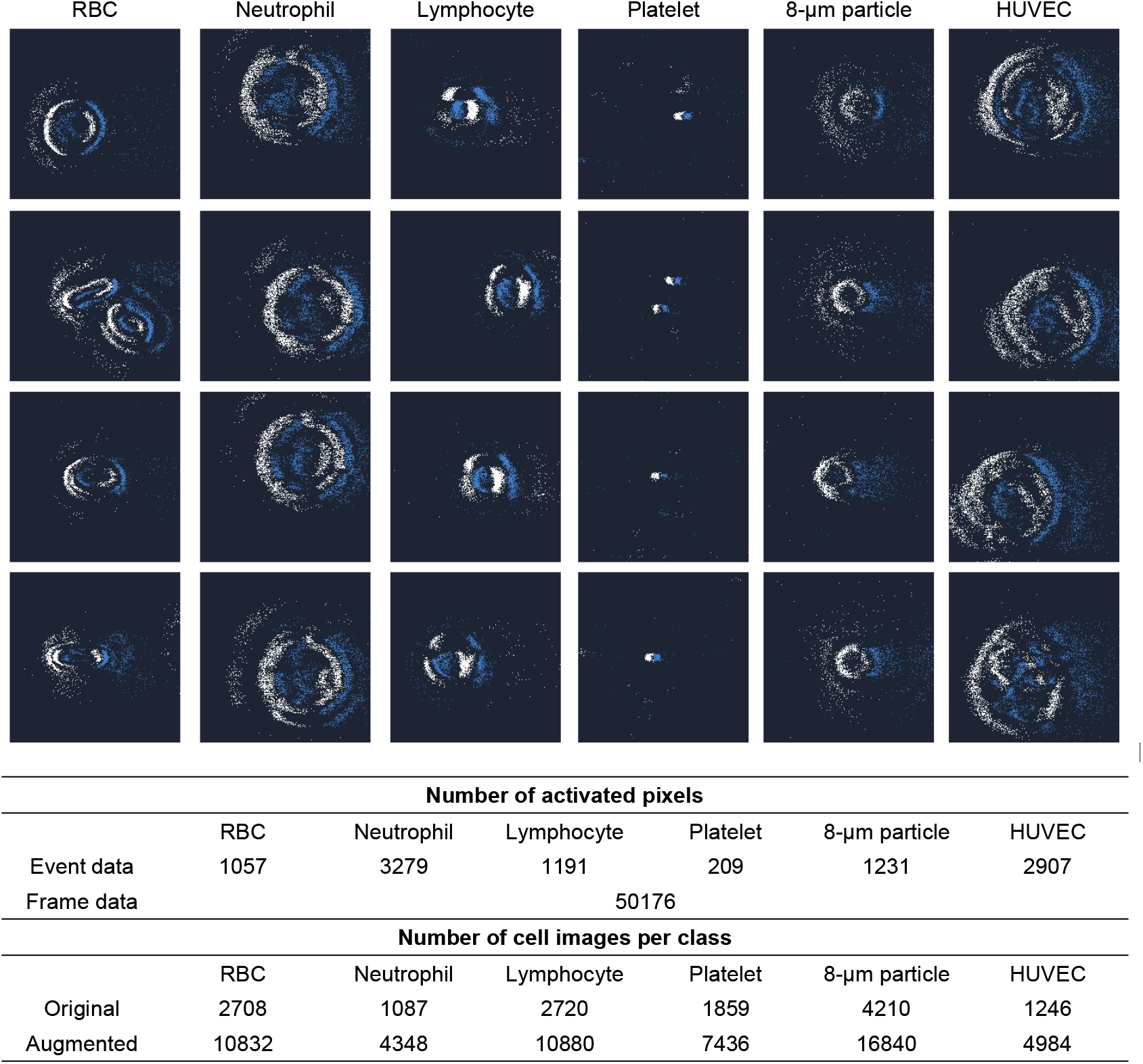
Neuromorphic data curation for human blood cells (RBCs, neutrophils, lymphocytes, platelets), 8-*μ*m particles and HUVECs illustrated in Metavision Studio dark blue theme representation. The average number of activated pixels per cell type was calculated in event data and compared with frame data. The original number of sample cells and the number of sample cells after data augmentation were reported.

The average number of activated pixels of each class under neuromorphic vision was recorded and compared with the frame data. Owing to the sparse activation in NVS, cells were registered only with correlated pixel addresses, where frame data contains a fixed pixel quantity regarding the default resolution, resulting in a significant difference in overall pixel recruitment. This difference can be exacerbated when observing particles of smaller sizes, as these objects only need a small proportion of pixels to characterise themselves. In this dataset, pixel recruitment in NVS has a 15–240 times reduction compared to the frame data.

The total number of collected cell images is also illustrated in Fig. 2. The discrepancy in cell images curated for each sample type can be due to the natural concentration difference in samples, loss during microfluidic delivery and the separation yields. Furthermore, the +/-polarity of NVS data is based on the light contrast changes. Objects travelling in different directions will lead to distinct changes in colour distribution. To encompass the cells travelling at different angles, we rotated each cell image at a 90-degree angle three times and generated a new cell image at each rotation. This augmentation technique can enhance the model to become invariant to orientation and expand the data volume.

### 2.3 Model Performance

In Fig. 3a, the learned features by the model were embedded into a Uniform Manifold Approximation and Projection (UMAP) to evaluate the high-dimensional feature extraction and feature correlation among classes. Most of the cell events were grouped into their respective clusters with a minor number of outliers resembling features across classes. Noticeably, the clusters of neutrophils and HUVECs were plotted relatively close together. This can be owing to the high resemblance of observable traits such as size and internal complexity. RBCs and 8 *μ*m particles were also distributed similarly. A confusion matrix was constructed based on the network’s prediction and ground truth in Fig. 3b. All predictive classes achieved a high ratio (> 0.94) matching with the true annotation. The classification for platelets, lymphocytes and 8 *μ*m particles accomplished outstanding scores of 1.00, 0.98 and 0.98. For ICS applications, the process time from generating an image to concluding a decision is critical to determine the throughput. A rank-ordered fraction of events based on processing time is shown in Fig. 3c with 98.3% events concluded within 1 ms. Working with EVS with SNN architecture allows direct inference with event data, rendering an extremely short interval for image construction and data transfer.

**Figure 3.**
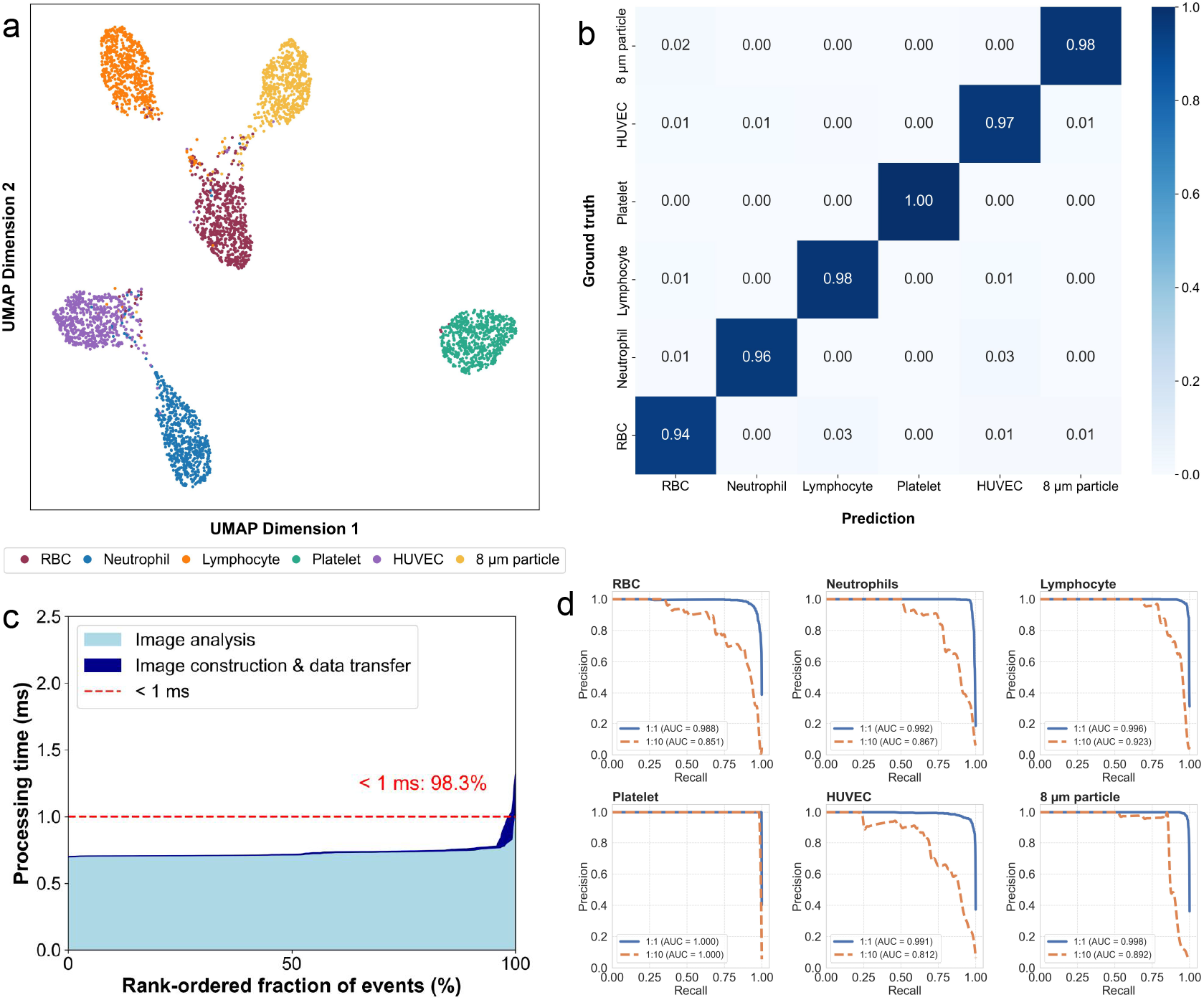
SNN model performance based on neuromorphic cell data. **a** Cell clusters in UMAP for revealing high-dimensional feature correlation in a 2D plot. **b** Confusion matrix based on each class’s prediction outcome and ground truth annotation. **c** Processing time for image construction and data transfer with class inference time in a hierarchical arrangement of events from lowest to highest processing time. **d** Precision-recall curve for validation trained with 1:1 balanced and 1:10 imbalanced datasets.

Furthermore, to accommodate the concentration variance among cells in a biological environment, for example, human blood consists of 40-45% RBCs, 1% white blood cells, 1% platelets or rare occurrences of endothelial cells in peripheral circulation,^18^ a balanced and imbalanced dataset were introduced to the network by maintaining a 1:1 ratio for targeted cells to total cells per class in the balanced case, and a 1:10 ratio in the imbalanced case. As shown in Fig. 3d, a precision-recall curve was plotted for balanced and imbalanced conditions of each class, with the score of the area under the curve (AUC) provided. The network exhibited a saturated curve with high precision and recall values in a 1:1 ratio, resulting in an average AUC score of 0.994. Even the RBC class, which had the lowest performance, achieved an AUC of 0.988, and the platelet class achieved a perfect score of 1.000. The model experienced a score drop with the imbalanced dataset but could still reach an average AUC of 0.891.

Herein, we explored the compatibility of an SNN network to predict cell events since SNNs can provide direct inference with event streams, unlike other networks, which require cumbersome preprocessing to transform events into frames. The performance of the CBAM-SNN-18, ResNet-18^19^ and EMS-Res-18^20^ were compared in Table 2. The ResNet-18 has achieved the highest performance in accuracy and F1 score of 99.82% with 0.75 ms processing time, whereas CBAM-SNN-18 achieved 97.17% accuracy and 97.16% and F1 score with 0.74 ms processing. Our model also reported a 1.77-fold and 39.15-fold reduction in inference memory usage and energy consumption. SNNs, owing to their binary activations, typically offer lower accuracy than conventional continuous processing networks in exchange for significantly improved efficiency in model size, energy usage, and suitability for real-time applications. In this work, we have trained a CBAM-SNN that has comparable performance to a classic CNN while outperforming memory and energy consumption efficiency. In addition, EMS-Res-18 is an SNN model that can also directly work with event files; our model achieved an increase in accuracy and F1 score under a similar architecture with a slight increase in computation resources. These characteristics indicate less network dependence on computational and hardware resources, suggesting an ease of miniaturised implementation in AI-enabled cytometers and sorters. A future endeavour to implement this architecture in neuromorphic hardware can lead to significant acceleration in latency and power gain.^21,22^

**Table 2:** CBAM-SNN-18, ResNet-18 and EMS-Res-18 performance comparison. F1-score is weighted. Processing time is the sum of image preprocessing and inference time per sample. Inference memory represents the VRAM consumption of the model per sample.

| | Accuracy (%) | F1-Score (%) | Processing Time (ms) | Memory (MB) | Parameter (M) | Energy Consumption (μJ) |
| --- | --- | --- | --- | --- | --- | --- |
| CBAM-SNN-18 | 97.17 | 97.16 | 0.74 | 93.46 | 6.32 | 215 |
| ResNet-18<sup>a</sup> | 99.82 | 99.82 | 0.75 | 165.17 | 11.69 | 8417 |
| EMS-Res-18<sup>b</sup> | 96.21 | 96.18 | 0.70 | 88.77 | 6.32 | 170 |

<sup>a</sup> ResNet-18 and <sup>b</sup> EMS-Res utilised in this work adopted the original architecture in He et al.<sup>19</sup> and Su et al.<sup>20</sup> trained on this dataset.

## 3 Method

### 3.1 Sample Preparation

Human blood samples were collected into EDTA tubes from healthy volunteers following written informed consent under the Sydney Local Health District ethics protocol X23-0300. To ensure each footage contains the respective cells while maintaining a sufficient sample volume for data labelling and training purposes, the samples were treated with individual approaches to bulk separate the blood cells. Firstly, the platelet containing the plasma layer was separated following centrifugation at a speed of 800 g for 15 minutes. The plasma-depleted blood, layered with Ficoll-Paque (GE Healthcare, Sweden) underwent gradient centrifugation at a speed of 500 g, with no brake for 30 minutes to separate layers of different compositions in the blood. Lymphocytes were retrieved from the interface layer. At the bottom layer of the sample, RBCs were retrieved via conventional pipetting technique and diluted by a factor of 100 with PBS to reduce the excessive concentration of RBCs in the original sample. A second aliquot of the bottom layer was treated with red blood cell lysis, to enrich for the remaining neutrophils.

Human umbilical vein endothelial cells (HUVECs, Lonza) were cultured in endothelial growth media (EGM-2, Lonza) at 37 °C and 5% CO_2_. Cells were passaged using TrypLE Express (Gibco) and used between passage 4 and passage 6 for experiments. To prepare cell suspension for microfluidic imaging experiments, dissociated HUVECS were resuspended in fresh warm media at a concentration of 6 × 10^5^ cells/mL and used within 30 minutes.

### 3.2 Microfluidic Imaging Platform

A standard soft-lithography protocol^26^ was used to fabricate a straight microfluidic channel (35 *μ*m-height, 100 *μ*m-width and 30 mm long) with polydimethylsiloxane (PDMS) (Sylgard® 184 Silicone Elastomer Kit, Dow Corning, UK). To ensure the sample cells were travelled in a common depth for in-focus imaging, the channel was designed with a limited height to narrow down the z-plane passage during the delivery. An event camera, Evaluation Kit 4 (EVK4, Prophesee, France) adopted in this experiment operated at 1280 × 720 resolution, 4.86 × 4.86 *μ*m pixel size with time resolution equivalent > 10, 000 frame per second (fps). The camera was integrated into an inverted microscope (IX73 Inverted Microscope, Olympus, Japan) to record the cell dynamics under 40 × magnification in a phase contrast mode. The magnification was chosen to enhance the captured features and reveal the unique internal complexity of the cells. Metavision Studio software provided by Prophesee was utilised during the recording to optimise the visual feedback at an accumulation time of 200 *μ*s.

For microfluidic delivery, the sample solution was loaded into a 1 mL syringe (Terumo Syringe, Terumo Corporation, Philippines) and actuated into the channel via a syringe pump (LEGATO 200 Syringe Pumps, KD Scientific Inc, USA) to induce a flow velocity of 4.78 mm/s. The flow velocity adopted in this experiment permits sufficient exposure time to the sensor while maintaining a relatively fast speed. A new channel was implemented after each delivery class to avoid sample contamination from the leftover particles.

### 3.3 Data Preprocessing

As the cells were captured at a resolution of 1280 × 720, which is not an ideal size for single-cell analysis and import for machine learning, the event data was first cropped into a 224 × 720 region of interest. A filtering technique that sets an event threshold of 300 was applied to eliminate the potential background noise and cell debris in the solution. These events were removed if the total event counts were under the threshold within the period. As the size of platelets is much smaller compared to the rest of the samples, a threshold of 50 was adopted for the platelet batch.

The cell events were then cropped into 224 × 224 using K-means Clustering to capture the samples at the centre of the images and data array. The event stream was divided into individual samples with a duration of 200 *μ*s to generate 2D images for annotation and tensors with dimensions (X, Y, P, T) for model training. Furthermore, outliers with radical morphology differences, blurry edges, and out-of-focus were manually discarded by the researcher, who had empirical experience working with EVS and targeted cell lines.

### 3.4 Model Development

Event data can be cumbersome to process with traditional neural networks regarding the unique event generation mechanism. Spiking neural networks are a natural match for implementation since they can be trained directly and exploit temporal information. It is also characterised by low latency, superior memory and energy efficiency. The forward propagation action in this network can be represented by Equ. 1:^27^

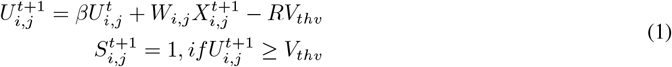

where 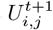 and 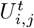 represented the membrane potential (*V*) in location (*i, j*),*W*_*i,j*_ is the weight for different input spikes, *S* is the input spikes (V), *β* is the learnable parameter controlling membrane potential decay, *R* is the parameter represent the reset mechanism.

Surrogate Gradient was adopted to train our SNN network directly. It replaced the infinite gradient with the arc-tan function in the backpropagation process to solve the dead neurons problem, which is shown in Equ 2.^27,28,29^

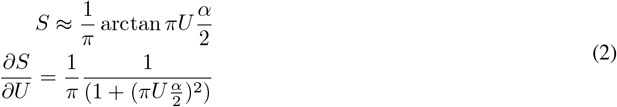

where *S* is the spikes this neuron produced (*V*), *U* is membrane potential(*V*), and *α* is a parameter that defaults to 2.

The model architecture of CBAM-SNN is demonstrated in Fig. 4a. As shown in Equ 1, the LIF layer does not account for information from nearby neurons because its behaviour depends only on the history of its own signals. Therefore, the convolutional layers in a Convolutional SNN are responsible for extracting spatial features. However, a few convolutional layers with small kernels in CSNN cannot sufficiently capture spatial features because of their limited receptive fields.^31^ Hence, to further enhance the network’s capability without introducing excessive parameters and unnecessary computations, a local attention block, CBAM as shown in Fig 4b, was implemented into the SNN-ResNet structure. This block is a widely known local attention mechanism that can help the network focus more on the important part of the input data.^32^ It has already been used in lots of lightweight networks due to its efficiency.^33,34,35^ The combined local attention mechanism is expressed as Equ. 3,

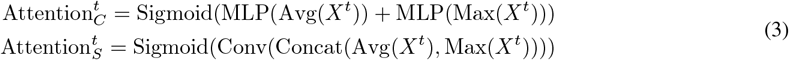

where *t* represents the current time step, *X* is the input spikes and MLP(·) is the shared multilayer perception, which has the same structure as SE-Block,^36^ Avg(·) is global average pooling, Max(·) is global Max pooling and Concat(·) represents the action concatenating two tensors in Channel dimension.

**Figure 4.**
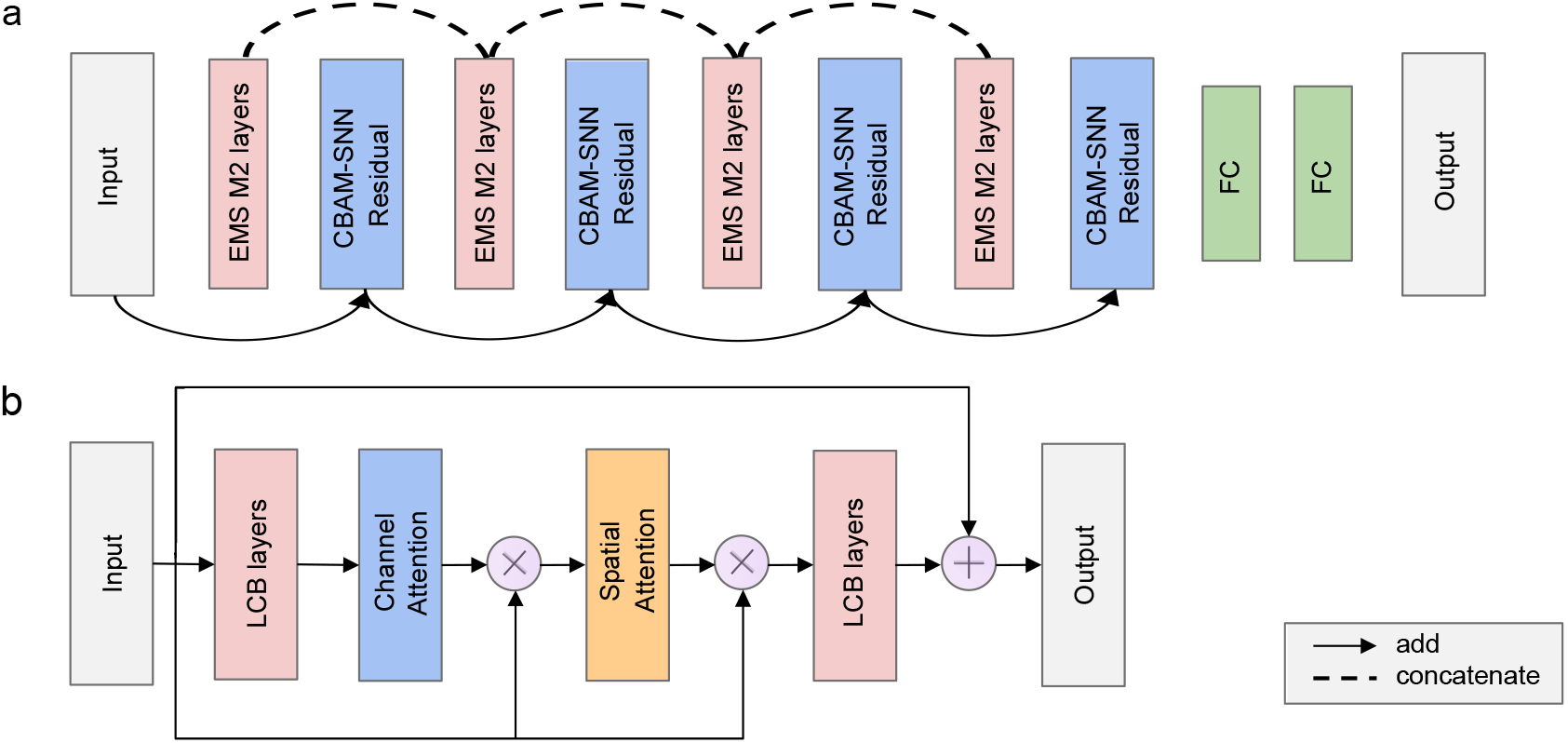
CBAM-SNN model overall architecture. **a** The general structure of the CBAM-SNN model. Each basic block has three layers: one downsampling energy-efficient membrane-shortcut (EMS) block and two SNN-Residual blocks. Two bottleneck^30^ LCB layers construct the SNN-Residual block. One local attention CBAM block would be inserted into two LCB layers to build up the CBAM-SNN Residual layer. **b** The structure of the CBAM-SNN Residual block. When spikes enter this block, they will first go through one LCB layer; channel-wise attention is calculated in Equ 3 and multiplied with the input spikes. The result of multiplying will act as the input of the spatial attention layer, repeat the previous operation, and then pass the result to the final LCB block.

All the blocks are built upon the fully spiked Energy-Efficient Residual Block, EMS block,^20^ which consists of two branches, one consisting of two continued LCB blocks constructed by Leaky-integrate-fire network, batch normalisation layer and convolution layer sequentially. The other branch is the residual branch, which consists of one Max-pooling layer used to transform the shape of the input data and avoid involving more MAC operation and one LCB block. In the second branch, the output from two components would be concatenated and act as the result of the residual branch. The final output of this layer is the sum of these two branches, which is a kind of full spike residual connection.

To avoid the possibility of under-fitting during the training,^37,38^ the skip connection was added to all the local attention blocks, and some residual connections were added to each down-sampling layer. These connections greatly enhance the performance of the model and render the training processes more stable.^30,39^ To further reduce the MAC operation, all skip and residual connections use the same structure as EMS-ResNet used in EMS-Module 1,^20^ which consists of Max-pooling and LCB blocks. Moreover, depth-separable convolution layers were implemented to replace the normal convolution layer to reduce the computation consumption.^40,41^All the training and validation were operated on one Linux platform, Ubuntu 22.04, with an Intel i5 13600KF CPU and Nvidia RTX 2080TI (22GB VRAM). The AdamW optimised all the models using the learning rate reduce scheduler, and the start learning rate was 0.0001.

## 4 Conclusion

NVS has been vastly adopted in the field of auto-driving, aerial drone vision and robotics, owing to its superior spatiotemporal resolution, data efficiency, low power consumption and high dynamic range.^42,43,44^ However, this architecture has not yet been fully recognised and utilised in biological cell research using cytometric approaches, especially for handling large-scale cell data with high dimensionality. In this work, we presented the first visualisation of human blood cells under neuromorphic vision, assessing the event quality for representing the spatial content of cells. To demonstrate the possibility of DL to substitute conventional manual gating strategy and fast inference for any large data-required or time-sensitive application such as ICS or IFC, we developed a hybrid model of CBAM-SNN that can be trained directly on raw event data with reduced computational needs while providing a highly accurate fast inference. This unique event-focused sensing architecture can also be an excellent candidate for monitoring cell trajectory, motion dynamics, transient behaviours and 3D cytometry in any time- and motion-critical applications.

For future endeavours, our dataset currently contains six classes of cells, which is still limited to be considered as generalised and more cells appear in greater similarity. Thus we will continue to build on this dataset to collect more cell lines or fine types of cells in exclusive contexts. Also, implementing the entire paradigm on neuromorphic hardware can enhance network performance by providing a faster inference and power gain.

Given that cells are typically cultured and studied in vast quantities, generating a full frame out of each cell can be even more data-expensive and challenging to examine by humans individually. Machine learning operates with convoluted algorithms to reveal meaningful data structure and requires a large data volume to train and conclude solid inference. Combining event-based sensing with DL can render a data-efficient, high throughput and fluorescence-sensitive paradigm to overcome the conventional trade-off in IFC and ICS, laying down a promising milestone for next-generation AI-assisted cytometry/sorting applications.

## 5 Ethic Statement

Human blood samples are obtained with full informed consent from healthy volunteers under the Sydney Local Health District ethics protocol X23-0300 in 2023. All procedures comply with institutional review board guidelines and respect participant privacy, confidentiality, and autonomy. Samples are used solely for research purposes, adhering to ethical guidelines for human biological materials.

## 6 Data Availability

All codes and data repositories are released under appropriate open-source licenses and are available via https://github.com/NeuroSyd/CBAM-SNN. Neuromorphic Cell Dataset can be accessed via https://doi.org/10.6084/m9.figshare.27788970.

## 7 Acknowledgement

SWC acknowledges funding from The University of Sydney’s Faculty of Engineering Scholarship.

## 8 Author Contribution

Z.Z. designed and conducted all the physical cytometry and biological experiments as well as the paper write-up. H.Y. developed the machine learning model to conduct cell classification tasks. J.L. and Z.Z performed data analysis, visualisation and initial platform design. S.W.C. and H.M.M. prepared biological samples for the experiment. J.K.E participated in the initial conceptualisation and continuously provided professional guidance on neuromorphic sensor and model development. D.V. and K.T.Y. designed and offered microfluidics expertise during the research. O.K. conceptualised the idea, initiated the project, conceptualised the experiments and edited the paper. O.K., D.V. and H.M.M. jointly provided primary supervision.

## 9 Supplementary Information

Sample supplementary video footage of human blood cells are available via https://doi.org/10.6084/m9. figshare.27789078. Additional data curation and model development information can be viewed in supplementary content.

